# Multinucleotide mutations cause false inferences of lineage-specific positive selection

**DOI:** 10.1101/165969

**Authors:** Aarti Venkat, Matthew W. Hahn, Joseph W. Thornton

**Affiliations:** Department of Human Genetics, University of Chicago, Chicago IL 60637, USA; Department of Biology and Department of Computer Science, Indiana University, Bloomington IN 47405, USA; Department of Ecology & Evolution, University of Chicago, Chicago IL 60637, USA

**Keywords:** adaptation, adaptive evolution, branch-site test, codon models, transversions

## Abstract

Phylogenetic tests of adaptive evolution, which infer positive selection from an excess of nonsynonymous changes, assume that nucleotide substitutions occur singly and independently. But recent research has shown that multiple errors at adjacent sites often occur in single events during DNA replication. These multinucleotide mutations (MNMs) are overwhelmingly likely to be nonsynonymous. We therefore evaluated whether phylogenetic tests of adaptive evolution, such as the widely used branch-site test, might misinterpret sequence patterns produced by MNMs as false support for positive selection. We explored two genome-wide datasets comprising thousands of coding alignments – one from mammals and one from flies – and found that codons with multiple differences (CMDs) account for virtually all the support for lineage-specific positive selection inferred by the branch-site test. Simulations under genome-wide, empirically derived conditions without positive selection show that realistic rates of MNMs cause a strong and systematic bias in the branch-site and related tests; the bias is sufficient to produce false positive inferences approximately as often as the branch-site test infers positive selection from the empirical data. Our analysis indicates that genes may often be inferred to be under positive selection simply because they stochastically accumulated one or a few MNMs. Because these tests do not reliably distinguish sequence patterns produced by authentic positive selection from those caused by neutral fixation of MNMs, many published inferences of adaptive evolution using these techniques may therefore be artifacts of model violation caused by unincorporated neutral mutational processes. We develop an alternative model that incorporates MNMs and may be helpful in reducing this bias.

## INTRODUCTION

Identifying genes that evolved under the influence of positive natural selection is a central goal in molecular evolutionary biology. During recent decades, likelihood-based phylogenetic methods have been developed to identify gene sequences that retain putative signatures of past positive selection ^1–10^. Perhaps the most widely used of these is the branch-site test (BST) of episodic selection, which allows positive selection to affect only some codons on one or a few specified branches of a phylogeny, and therefore has relatively high power compared to methods that detect selection across an entire sequence or an entire phylogenetic tree ^5,6,11^. The BST has been the basis for published claims of lineage-specific adaptive evolution in many thousands of individual genes ^12–16^.

The BST and related methods use a likelihood ratio test to compare how well two mixture models of sequence evolution on a phylogeny fit an alignment of coding sequence data. The null model constrains all codons to evolve with rates of nonsynonymous substitution (*d*_N_) less than or equal to the rate of synonymous substitution (*d*_S_), as expected under purifying selection and drift. In the positive selection model, some sites are allowed to have *d*_N_>*d*_S_ on a branch or branches of interest. If the increase in likelihood of this model given the data is greater than expected due to chance alone, the null model is rejected and adaptive evolution is inferred. The BST has been shown to be conservative, with a low rate of false positive inferences, when sequences are generated under an evolutionary process corresponding to the null model ^6,11^. It is widely appreciated that likelihood ratio tests can become biased if the underlying probabilistic model is incorrect ^17^. The effect on the BST of a few forms of model violation—such as an unequal distribution of selective effects among sites, positive selection on non-foreground lineages, high sequence divergence, and non-allelic gene conversion—have been previously studied ^18–22^ and the test has been found to be reasonably robust to most but not all forms of violation examined ^6,23,24^.

Recent research in molecular genetics and genomics suggests a potentially important phenomenon that has not been incorporated into models used in tests of positive selection: the propensity of DNA polymerases to produce mutations at neighboring sites. All implementations of the BST and other likelihood-based tests of adaptive evolution use models in which mutations occur only at individual nucleotide sites and are fixed singly and independently. Codons with multiple differences between them can be interconverted only by serial single-nucleotide substitutions, the probability of which is the product of the probabilities of each independent event. Recent molecular studies have shown, however, that mutations affecting adjacent nucleotide sites often occur during replication, apparently because certain DNA microstructures recruit error-prone polymerases that lack proofreading activity and therefore make multiple errors close together ^25–33^. Consistent with these mechanisms, genetic studies of human trios and mutation-accumulation experiments in laboratory organisms indicate that *de novo* mutations occur in tandem or at nearby sites more frequently than expected if each occurred independently ^25,32–36^, and these multinucleotide mutations (MNMs) are enriched in transversions ^35,37,39^. The precise frequency at which MNMs occur is difficult to estimate, but a recent compilation of genetic studies in humans concluded that about 0.4% of mutations, polymorphisms, and substitutions are at directly adjacent sites (counting each tandem pair as one event) ^34^. In *Drosophila melanogaster* genomes, analysis of rare polymorphisms and mutation-accumulation experiments estimated that 1.3% of all mutations are at adjacent sites ^38^. Although the methods and data sources in these studies differ, these findings suggest that tandem MNMs probably account for on the order of 1% of mutational events.

We hypothesized that these mutational processes might lead to false signatures of positive selection in the BST. Because of the structure of the genetic code, virtually all MNMs in coding sequences are nonsynonymous, and most would comprise multiple nonsynonymous nucleotide changes if they were to occur by single nucleotide steps (**Supplementary Table 1**). The enrichment of transversions in MNMs further increases the propensity for MNMs to produce nonsynonymous changes, because transversions are more likely than transitions to be nonsynonymous. MNMs are therefore likely to produce codons with multiple differences (CMDs) that contain an apparent excess of nonsynonymous substitutions. When these CMDs are assessed using a method that treats all substitutions as independent events, a model that allows *d*_N_ to exceed *d*_S_ at some sites may have a higher likelihood than one that restricts *d*_N_/*d*_S_ to values ≤ 1. Further, the assumption that all mutations have the same transversion-transition rate might exacerbate the tendency to misinterpret MNM-produced nonsynonymous changes as evidence for positive selection. Of course, CMDs can also be driven to fixation by positive selection ^11,40–42^—and the same is true of transversion-rich substitutions—but these considerations suggest that failing to incorporate MNMs in likelihood models might make tests of adaptive evolution susceptible to false positive inferences. The BST and other lineage-specific tests might be particularly sensitive to this problem because they seek signatures of positive selection acting on small numbers of codons on one or a few specified branches of the tree ^43^. Simulation studies suggest that MNMs may elevate false positive rates in some selection tests ^44^, but there has been no comprehensive analysis of the effect of MNMs, particularly on the branch-site test or under realistic, genome-scale conditions.

## RESULTS

To understand the effect of MNMs on the accuracy of the branch-site test, we analyzed in detail two previously published genome-wide datasets, which represent classic examples of the application of the test ^13,15,45^. The mammalian dataset consists of coding sequences of 16,541 genes from six eutherian mammals; we retained for analysis only the 6,868 genes with complete species coverage. The fly dataset consists of 8,564 genes from six species in the melanogaster subgroup clade, all of which had complete coverage (**Supplementary Fig. 1**). The fly genes have higher sequence divergence than those in the mammalian dataset, allowing us to examine the performance of the BST under different evolutionary conditions.

We used the classic BST to identify genes putatively under positive selection (P<0.05) on the human lineage in the mammalian dataset and on each of the six terminal lineages in flies. 82 genes in humans and 3,938 in flies yielded significant tests (**Supplementary Table 2**). To facilitate robust further analysis of CMDs, we filtered out genes in which CMDs occurr at sites with indels or in which the ancestral states of CMDs are reconstructed differently between the null and positive selection models; we also applied a multiple testing correction (FDR <0.20). In flies, 443 genes were retained after these steps. Thirty human genes passed the CMD alignment and reconstruction filter, but none met the FDR threshold, consistent with previous analyses of these data; ^15^ nevertheless, we included the 30 initially significant human genes because this lineage is the object of intense interest and because its short length contrasts with the fly branches. These two groups constitute the “BST-significant” sets of genes in flies and humans.

### CMDs provide virtually all support for positive selection

We sought to determine how much of the evidence for positive selection comes from CMDs. We first observed that CMDs were dramatically enriched in BST-significant genes compared to non-BST-significant genes (**Fig. 1a**). In humans, BST-significant genes contain one CMD on average, while BST-nonsignificant genes contain none (**Supplementary Fig. 2**). The pattern is similar but less extreme in flies, with the average number of CMDs per BST-significant gene greater than that in non-significant genes (**Supplementary Fig. 2**). When CMD-containing codons are excluded from the alignments, the vast majority of genes that were BST-significant lose their signature of selection in both datasets (**Fig. 1b)**.

**Figure 1.**
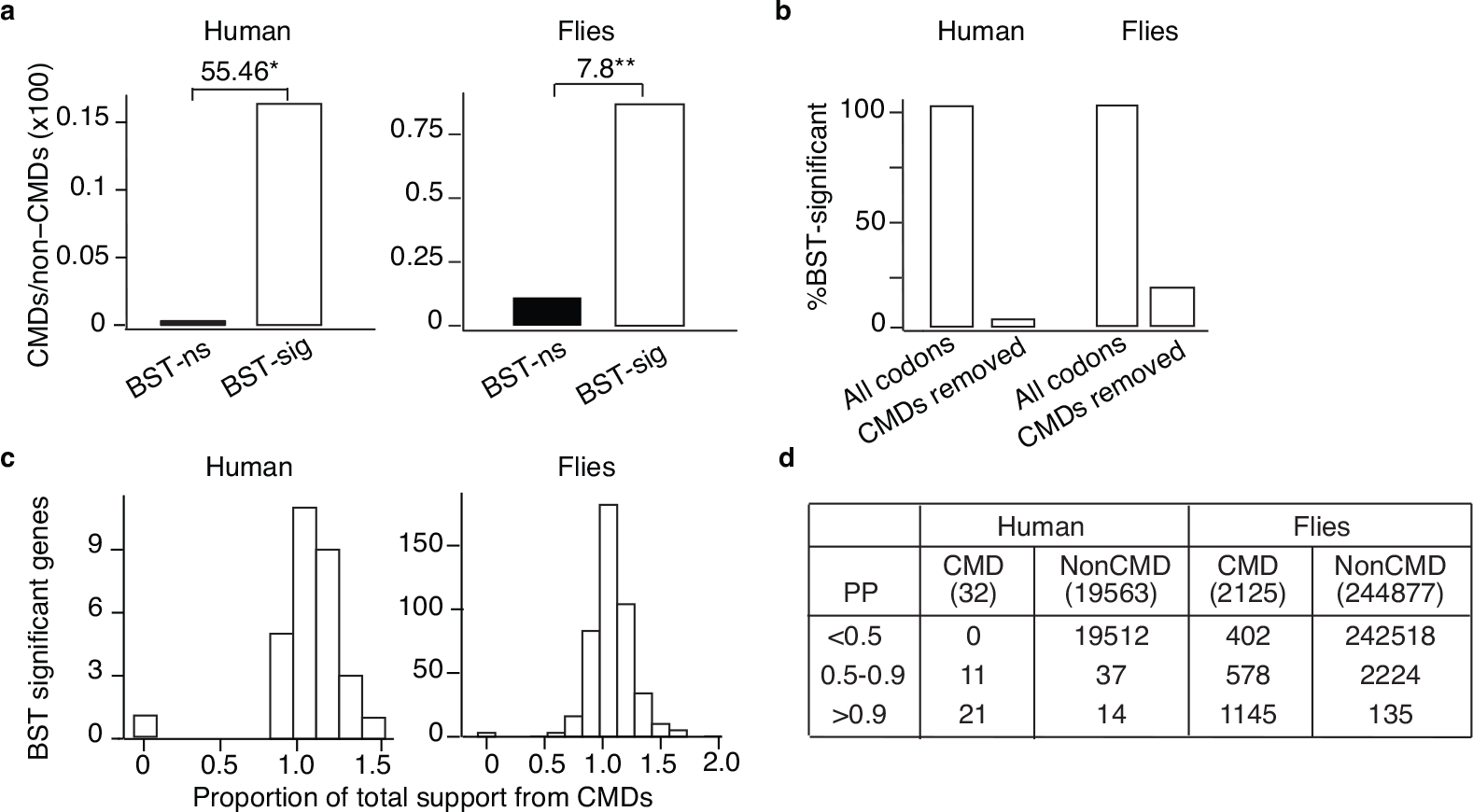
Codons with multiple nucleotide differences (CMDs) drive branch-site signatures of selection. **(a)** CMDs are enriched in genes with a signature of positive selection. Codons were classified by the number of nucleotide differences between the ancestral and terminal states on branches tested for positive selection. CMDs have ≥2 differences; non-CMDs have ≤1 difference. The CMD/non-CMD ratio is shown for genes with a significant signature of selection in the BST (BST-sig) and those without (BST-ns). Fold-enrichment is shown as the odds ratio. *, P=4e-4 by χ^2^ test; **, P=1e-41 by Fisher’s exact test. **(b)** Percentage of genes that retain a signature of positive selection when CMDs are excluded from the branch-site test analysis. **(c)** Distribution across BST-significant genes of the proportion of total support for the positive selection model that is provided by CMDs. Total support is the difference in log-likelihood between the positive selection and null models, summed over all codons in the alignment. Support from CMDs is summed over codons with multiple differences. The proportion of support from CMDs can be greater than 1 if the log-likelihood difference between models is negative at non-CMDs. **(d)** Most codons classified as positively selected are CMDs. The number of CMDs and non-CMDs in BST-significant genes are shown according to their Bayes Empirical Bayes posterior probability (PP) of being in the positively selected class.

We next calculated the fraction of statistical support for positive selection that comes from CMDs. The total support for positive selection in an alignment is defined as the difference between the log-likelihood of the positive selection model and that of the null model, summed across all codons in the alignment. The fraction of support from CMDs is the support from CMD-containing codons divided by the total support across the entire alignment. CMDs account for >95% of the support for positive selection in virtually all BST-significant genes in both datasets; in about 70% of genes, CMDs provide all the support (**Fig. 1c)**.

Finally, we examined the BST’s *a posteriori* identification of sites under positive selection. We found that CMDs were far more likely to be classified as positively selected than non-CMDs. Among genes that were BST-significant on the human lineage, every CMD was inferred to be under positive selection using a Bayes Empirical Bayes posterior probability (PP) cutoff > 0.5. Using a more stringent cutoff of PP>0.9, 66 percent of CMDs were classified as positively selected, compared to 0.07% of non-CMDs. In the fly dataset, CMDs accounted for 90% of codons with BEB>0.9, although they represent less than 1% of all codons (**Fig. 1d**).

CMDs are therefore the primary drivers of the signature of selection identified in the BST. A single CMD provides sufficient statistical support to yield a signature of positive selection on the human lineage, and only a few CMDs in a gene are enough to do the same in flies.

### Incorporating MNMs eliminates the signature of positive selection in many genes

CMDs might be enriched in BST-positive genes because of an MNM-induced bias or because they were fixed by positive selection. To incorporate both neutral and selection-driven fixation of MNMs into a BST framework, we developed a codon model in which double-nucleotide changes are allowed, with the parameter δ serving as a multiplier that modifies the rate of each double-nucleotide substitution relative to single-nucleotide substitutions. We implemented a version of the BST (BS+MNM) that is identical to the classic version, except that both the null and positive selection models allow MNMs. Simulations under conditions derived from a sample of genes in the mammalian dataset show that the method estimates the parameters used to generate the sequences with reasonable accuracy (**Supplementary Fig. 3**).

We first fit the BS+MNM null model to all alignments in the mammalian and fly datasets. The average estimate of δ across all genes was 0.026 in mammals and 0.062 in flies, with δ in both cases about twice as high in the subset of BST-significant genes as in BST-nonsignificant genes (**Fig. 2a)**. Using a likelihood-ratio test, we found significant support for the BS+MNM null model (compared to the classic BST null model) in 22% of human genes and >50% of fly genes (**Supplementary Table 3**); simulations without MNMs showed that this comparison has a very low false-positive rate (**Supplementary Table 4).**

**Figure 2.**
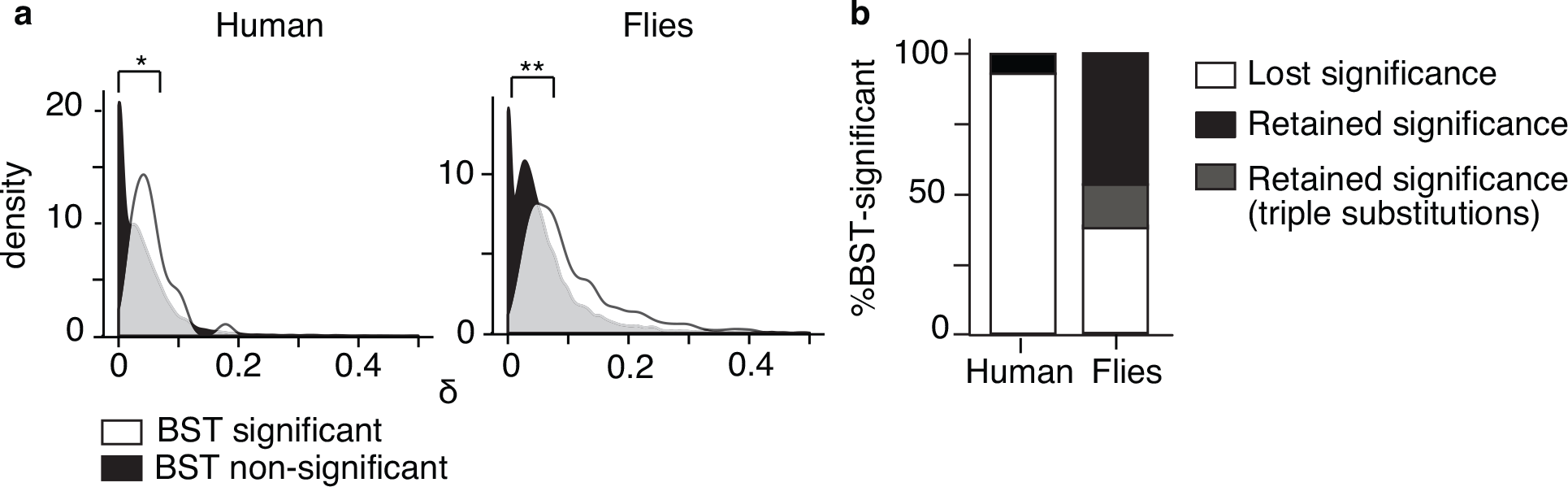
Incorporating MNMs into the branch-site model eliminates the signature of positive selection in many genes. The mammalian and fly datasets were reanalyzed using a version of the BST that allows MNMs (BS+MNM) by including a parameter δ, a multiplier on the rate of each double substitution relative to single substitutions. **(a)** The distribution of ML estimates of δ across genes with (white) and without (black) a significant result in the classic BST is shown for empirical alignments. Median estimates of δ in BST-significant and BST-nonsignificant genes are 0.047 and 0.026 in humans, respectively, and 0.107 and 0.062 in flies. *, P=6.7e-4; **, P=1e-8 by Mann-Whitney U Test. **(b)** Proportion of genes with a significant result in the BST that lose or retain that signature using the BS+MNM test. Genes that remain significant but contain CMDs with three differences, which are not incorporated into BS+MNM, are also shown.

We then used this BS+MNM test to evaluate the empirical sequences for positive selection. We found that 96% of the BST-significant genes on the human lineage lost significance in the BS+MNM test (**Figs. 2b, Supplementary Table 5**). In flies, 38% of the BST-significant genes lost significance; a substantial fraction of those that retained significance were enriched in triple substitutions, a process not accounted for in our model (**Figs. 2b, Supplementary Table 5**).

### MNMs cause false positive inferences on a genome-wide scale

That the BS+MNM test eliminates the signature of positive selection from many genes could have several causes, including: 1) the more complex BS+MNM model may have reduced power to identify authentic positive selection compared to the BST, 2) incorporating MNMs may ameliorate a bias towards false positive inference in the classic BST that is caused by MNMs, and 3) the additional δ parameter in the BS+MNM test may allow it to incorporate other forms of sequence complexity, potentially ameliorating a bias caused by other model violations.

We addressed these possibilities in two ways. First, we performed power analyses of the BS+MNM test using simulations in which positive selection is present in the generating model. We simulated sequence data on the mammalian and fly phylogenies using genome-wide averages for all parameters of the BST positive selection model, but we varied the strength of positive selection (ω_2_) and the proportion of sites under positive selection. We then applied the BS+MNM test to these data and found that it can reliably detect strong positive selection (ω_2_ > 20) when it affects more than 10% of sites in a typical gene, or moderate positive selection (10 < ω_2_ < 20) that affects a larger fraction of sites (**Supplementary Fig. 4a**). Under parameters derived from both datasets, the test’s power is similar to that of the classic BST, with slight reductions under only a few conditions on the fly lineage. Thus, although some genes may have lost their signature of selection because of reduced power in the BS+MNM test, it appears unlikely that a difference in power is the primary cause of the dramatic reduction in the number of positive results when the test is used.

Second, we used simulations under null conditions to directly evaluate the frequency of false positive inferences by the classic BST when sequences are generated with realistic rates of multinucleotide mutation. For every gene in the mammalian and fly datasets, we simulated sequence evolution under the null BS+MNM model without positive selection using parameters derived from the alignments, including δ. These parameters generate sequences with an observed frequency of tandem substitutions of 1.6% in humans and 3.2% in the *D. melanogaster* lineage in flies, similar to or slightly higher than the observed frequencies in the empirical datasets (1.3% and 1.6%, respectively), presumably because the BS+MNM model captures some but not all aspects of real sequence evolution (**Supplemental Table 6**) ^34, 38^.

We then analyzed these positive-selection-free simulated data using the classic BST. In both humans and flies, the number of genes with significant results—all of which are false positive inferences—was greater than the number of genes that the BST had concluded were under positive selection using the empirical data (**Fig. 3a).** In flies, almost 9 percent of tests were false positives (P<0.05), despite the conservative approach the method uses to calculate P-values ^6,11^, compared to just 1 percent under control simulations without MNMs. Further, more than 1,700 of these false positive tests survived FDR adjustment, compared to just 4 in the control simulations (**Supplementary Table 2)**. In humans, the fraction of false positive inferences is lower, consistent with the test’s reduced power in this dataset, but still about three times greater than in the control simulations.

**Figure 3.**
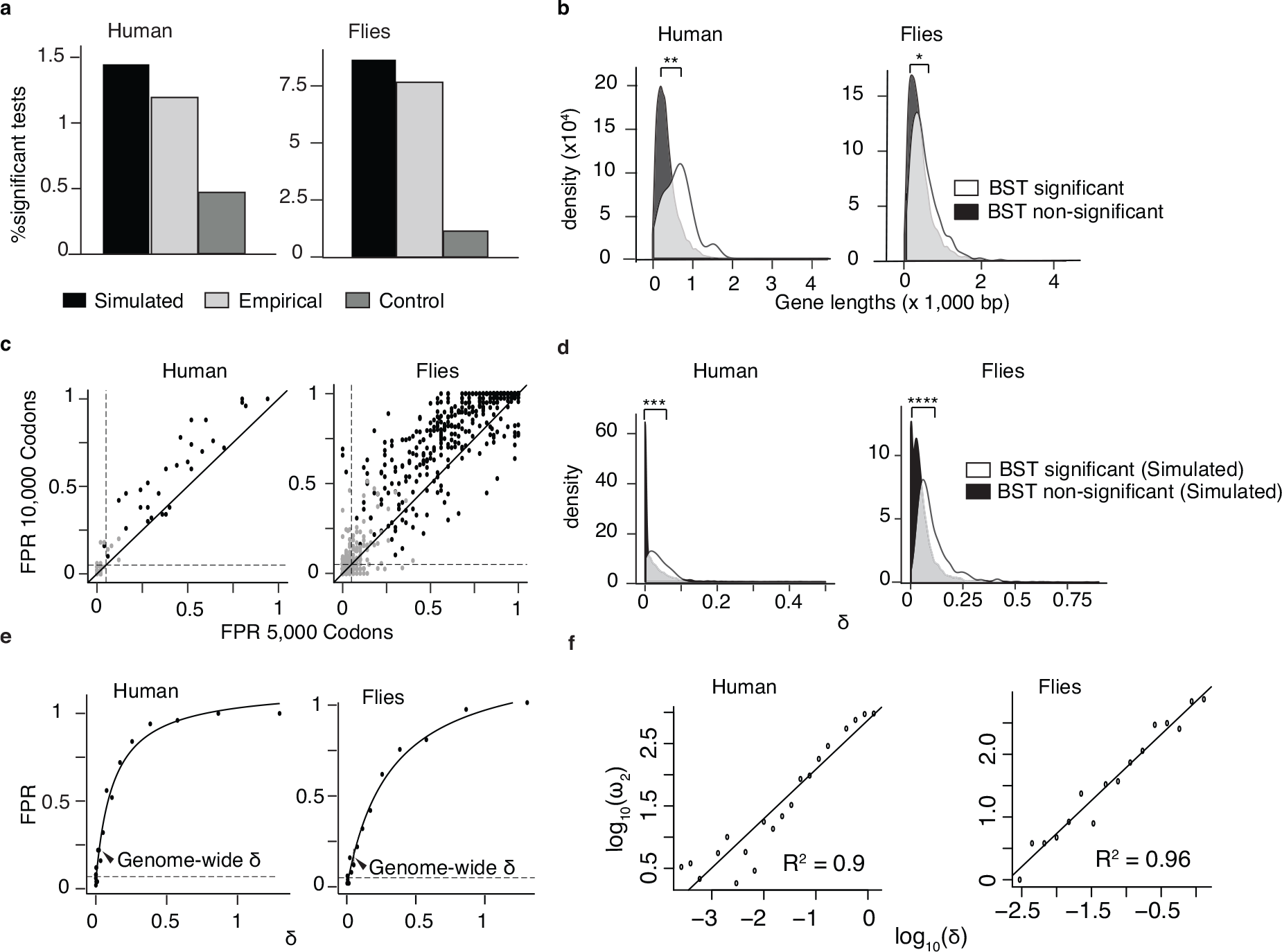
MNMs cause a strong bias in the branch-site test under realistic conditions. For each gene in the mammalian and fly datasets, the parameters of the BS+MNM null model were estimated by maximum likelihood. We then simulated sequence evolution under each gene’s inferred null parameters and used the classic BST on the simulated alignments to test for positive selection on the human and terminal fly lineages. **(a)** The fraction of all tests that are BST-significant (P<0.05) is shown for the data simulated under the BS+MNM null model, the original empirical sequence alignments, and a control dataset simulated with δ = 0. Each gene’s length in the simulation was identical to its empirical length. **(b)** BST-significant genes are longer than BST non-significant genes. The probability density of gene lengths in the two categories is shown for the empirical mammalian and fly datasets. Median lengths in BS-significant and non-significant genes, respectively, were 642 and 343 bp in humans; in flies, 448 and 399 bp. The difference between the two distributions was evaluated using a Mann-Whitney U test. *, P=8e-4; **, P=8e-5; **(c)** Systematic bias in the BST. For each gene with a significant result in the BST using the empirical data, we simulated 50 replicates using the BS+MNM null model and the ML parameter estimates for that gene at lengths of 5,000 and 10,000 codons; these data were then analyzed using the BST. The false positive rate (FPR) for any gene’s simulation (black points) is the proportion of replicates with P<0.05. Gray points show FPR for control simulations with δ = 0. Dashed lines, FPR of 0.05. The solid diagonal line has a slope of 1. **(d)** The distribution of ML estimates of δ across genes with (white) and without (black) a signature of positive selection in the classic BST is shown for data simulated under the BS+MNM null model. Median δ in BST-significant and BST-nonsignificant genes = 0.03 and 0.0009 in humans, 0.04 and 0.08 in flies. Difference between the distributions was evaluated using a Mann-Whitney U Test. ***, P=1e-12; ****, P=1e-199. **(e)** Increasing the MNM rate increases bias in the BST. Sequences 5,000 codons long were simulated using the BS+MNM model and the median value of each model parameter and branch length across all genes in each dataset, but δ was allowed to vary. The rate of false positives (P<0.05) in 50 replicates at each value of δ is shown. Solid line, hyperbolic fit to the data; dotted line, FPR level of 5%. Arrowhead, median δ across all genes. **(f)** Relationship between δ and inferred ω_2_. Sequences simulated in (e) were used to infer the ω_2_ estimated by BST under the positive selection model, and the relationship plotted on a log-log scale. The best-fit linear regression line is shown along with the coefficient of determination.

These false inferences are caused primarily by MNM-induced bias, because simulating data under identical control conditions without MNMs (δ = 0) produced few positive tests. All other parameters were identical between the generating model and analysis models, so other forms of model violation do not contribute to the bias observed in the simulation experiments. Taken together, these findings indicate that MNMs under realistic evolutionary conditions produce a strong and widespread bias in the BST toward false inferences of positive selection. This bias is strong enough to cause the BST to make false inferences of positive selection at about the same rate as it infers selection in the real genomes of humans and flies. In the simulations, every positive result is false; in the tests of real sequences, the fraction of positive results that are true is unknown.

### Systematic bias caused by chance MNMs in longer genes

We next sought to identify the causal factors that determine whether a gene yields a false positive result in the BST because of MNM-induced bias. Most genes are only several hundred codons long, and only a few percent of mutations are MNMs, so on phylogenetic branches of short to moderate length many genes will contain no CMDs caused by multinucleotide mutations. The hypothesis that neutral fixation of MNMs contributes to inferences of positive selection in the BST predicts that a gene’s propensity to produce a BST-significant result should depend on factors that increase the probability it will contain one or more fixed MNMs by chance, including its length and the gene-specific rate at which MNMs occur within it.

We first tested for an effect of gene length on the results of the branch-site test. As predicted, we observed that BST-significant empirical genes were on average 100 and 16 codons longer than non-significant genes in the human and fly empirical datasets, respectively (**Fig. 3b**). The relationship between length and propensity to yield a BST positive result could arise because genes that present a larger “target” are more likely to undergo MNMs than shorter genes; alternatively, longer genes, by including more sites for analysis, might increase the power of the BST to detect authentic positive selection. However, in genome-wide simulations under the null model with no positive selection (but with δ>0), genes with false positive BSTs are longer than the non-significant genes by an average of 26 and 31 codons using the human and fly parameters, respectively (**Supplementary Fig. 5**). This result cannot be attributed to increased power to detect true positive selection and supports the conclusion that mutational target size contributes to a gene’s propensity to manifest MNM-induced bias by chance alone.

To directly test the causal relationship between sequence length and false-positive bias in the BST, we simulated sequence evolution at increasing sequence lengths, using evolutionary parameters derived from each of the BST-significant genes in the mammalian and fly datasets. For each gene’s parameters, we simulated 50 replicate alignments under the BS+MNM null model and then analyzed them using the classic BST (**Supplementary Fig. 6a**). The false positive rate for any gene’s simulations is defined as the fraction of replicates with a significant LRT in the classic BST, using a P-value cutoff of 0.05. When sequences 5,000 codons long were simulated, 96% of BST-significant genes had an unacceptable FPR (>0.05), with a median FPR of 0.39: increasing sequence length to 10,000 codons exacerbated the bias, with 100% of genes yielding an unacceptable FPR and a median FPR of 0.56 (**Fig. 3c**). In flies, a similar pattern was evident, and the false positive rates were even higher (median FPR=0.74 and 0.90 at 5,000 and 10,000 codons, respectively). Control simulations under identical conditions but with δ=0 led to very low FPRs (median 0.02 to 0.03 in both datasets), even with very long sequences (grey dots in **Fig. 3c**). A similar systematic and length-dependent bias also resulted when sequences were simulated under gene-specific conditions, but with δ fixed to its average across the thousands of BST-nonsignificant genes in each dataset (**Supplementary Fig. 6b**). Although the sequence lengths tested are longer than most real genes, these experiments directly establish that a gene’s probability of returning a significant BST result in the absence of positive selection is directly related to the target size it presents for chance fixation of MNMs.

We next evaluated whether the gene-specific rate of multinucleotide mutation affects a gene’s propensity to yield a positive result in the BST. As predicted, we observed that BST-significant genes in the empirical datasets had higher estimated δ than nonsignificant genes **(Fig. 2a**). Genes producing false positive results in the genome-wide null simulations under empirical conditions also tended to have higher δ (**Fig 3d**); this result that cannot be attributed to the possibility that δ might be fitting CMDs fixed by positive selection, because positive selection was absent from the generating model.

To directly test the effect of the neutral MNM substitution rate on the BST, we simulated sequences 5,000 codons long under the null BS+MNM model, with a variable δ and all other parameters fixed to their averages across all genes. We found that increasing δ led to a monotonic increase in the frequency of false positive inferences. The FPR was >0.05 when δ was only 0.001 and 0.013 on the human and fly lineages, respectively. When δ was equal to its genome-wide average (0.026 and 0.062 in mammals and flies), false positive inferences occurred at rates of 22 and 17 percent, respectively (**Fig. 3e**). As δ increased, so too did the inferred value of the parameter ω_2_, which represents the inferred intensity of positive selection in the model (**Fig. 3f**).

Typical evolutionary conditions are therefore sufficient to cause a strong and systemic bias in the BST. MNMs are rare, however, so longer genes and those with higher rates of multinucleotide mutation are more likely to undergo this process and manifest the bias. This view is further supported by the fact that fewer genes are BST-positive on the human branch – which is so short that substitutions of any type are rare, and MNMs even more so – than on the fly phylogeny, where branches are longer, more CMDs are apparent, and hundreds of genes have BST signatures of selection. Taken together, these findings suggest that although some genes with BST-significant results in empirical datasets could have evolved adaptively, many may simply be those that happened to fix multinucleotide substitutions by chance alone.

### Transversion-enrichment in CMDs exacerbates bias in the branch-site test

MNMs tend to produce more transversions than classical single-site mutational processes, so if CMDs are produced by MNMs, they should be transversion-rich ^35, 37, 39^. As predicted, we found that transversion:transition ratio is elevated in CMDs relative to that in non-CMDs by factors of three and two in mammals and flies, respectively (**Fig. 4a**). In the subset of BST-significant genes, CMDs have an even more elevated transversion:transition ratio, as expected if transversion-rich MNMs bias the test (**Fig. 4a**). These data are consistent with the hypothesis that a transversion-rich MNM process produced many of the CMDs in BST-significant genes, but it is also possible that positive selection could have enriched for transversions.

**Figure 4.**
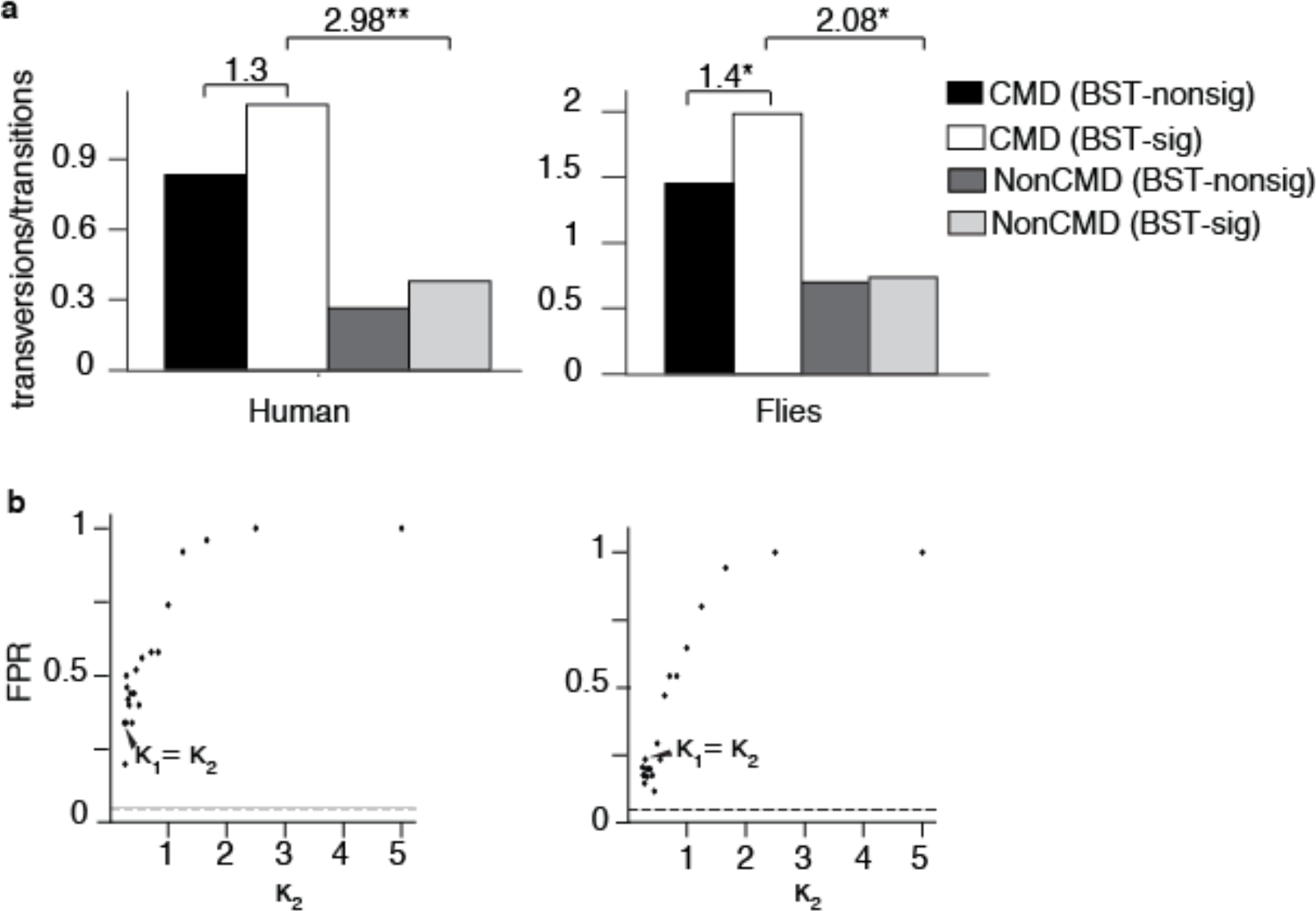
Transversion-enrichment in CMDs biases the BST. **(a)** The ratio of transversions:transitions observed in CMDs and in non-CMDs is shown for BST-significant and BST-nonsignificant genes. Fold-enrichment is shown as the odds ratio. *, P=5e-4; **, P=3e-25 by Fisher’s exact test. **(b)** Increasing the transversion rate in MNMs increases bias of the BST. Sequences 10,000 codons long were simulated using an elaboration of the BS+MNM model that allows MNMs to have a transversion:transition rate (κ_2_) different from that in single-nucleotide substitutions (κ_1_). 50 replicate alignments were simulated under the null model using the average value of every model parameter and branch length across all genes in each dataset, except κ_**2**_ was allowed to vary. The rate of false positives (P<0.05) at each value of κ_**2**_ is shown. Arrowheads show the false positive rate when sequences were simulated with κ_2_ equal to κ_1_. Dotted line, FPR of 5%.

To test whether transversion-enrichment in MNMs exacerbates the BST’s bias, we developed an elaboration of the BS+MNM model in which an additional parameter allows MNMs to have a different transversion:transition rate ratio (κ_**2**_) than single-site substitutions do (κ_**1**_). We estimated the maximum likelihood estimates of the model’s parameters for every gene in the mammalian and fly datasets and simulated sequences using genome-wide median values for all model parameters and branch lengths, except for κ_**2**_, which we varied. Sequences 10,000 codons long were used, because simulating shorter sequences resulted in a high variance in the realized transversion:transition ratio. We analyzed these data using the classic BST and calculated the fraction of replicates in which positive selection was inferred. We found that increasing κ_**2**_ caused a rapid and monotonic increase in the false positive rate, indicating that transversion enrichment in MNMs exacerbates the test’s bias. The effect is strong: when κ_2_/κ_1_ is increased from 1 to 2, the FPR approximately doubles (**Fig. 4b**). Thus, realistic rates of MNM generation and transversion-enrichment together cause an even stronger bias in the BST than MNMs alone. This result cannot be accounted for by positive selection driving fixation of transversions, because no positive selection was present in the simulations.

### MNMs affect a newer test of positive selection

In recent years, newer likelihood-based methods have been introduced to test for episodic site-specific positive selection^2,3,7^. All these methods are based on models of sequence evolution that, like the BST, do not allow MNMs but instead model CMDs as the result of serial site-specific substitutions. We therefore hypothesized that these methods might also be biased by MNMs. We chose a more recent method, BUSTED ^2^, which was developed primarily to test for episodic site-specific selection events across an entire tree. We tested its performance on alignments 5,000 codons long that were simulated using the BS+MNM null model and parameters estimated from the BST-significant gene alignments in humans and flies. To test for MNM-induced bias, we compared results when δ was assigned to three different values: zero, its average across all alignments in the mammalian or fly datasets, or its gene-specific value in each of the BST-significant genes (**Supplementary Fig. 6a**).

We found that BUSTED was also sensitive to MNM-induced bias. When δ=0, virtually no genes’ parameters led to frequent false positive inferences, with a median FPR <0.03 across genes **(Fig. 5)**. But when δ was assigned to its empirically estimated gene-specific value, the parameters from every gene in humans and the majority in flies yielded false positive rates >0.05, with median FPRs of 0.29 and 0.5, respectively (**Fig. 5**). Frequent false positive inferences were evident when sequences were simulated using genome-wide average estimates of δ, as well.

**Figure 5.**
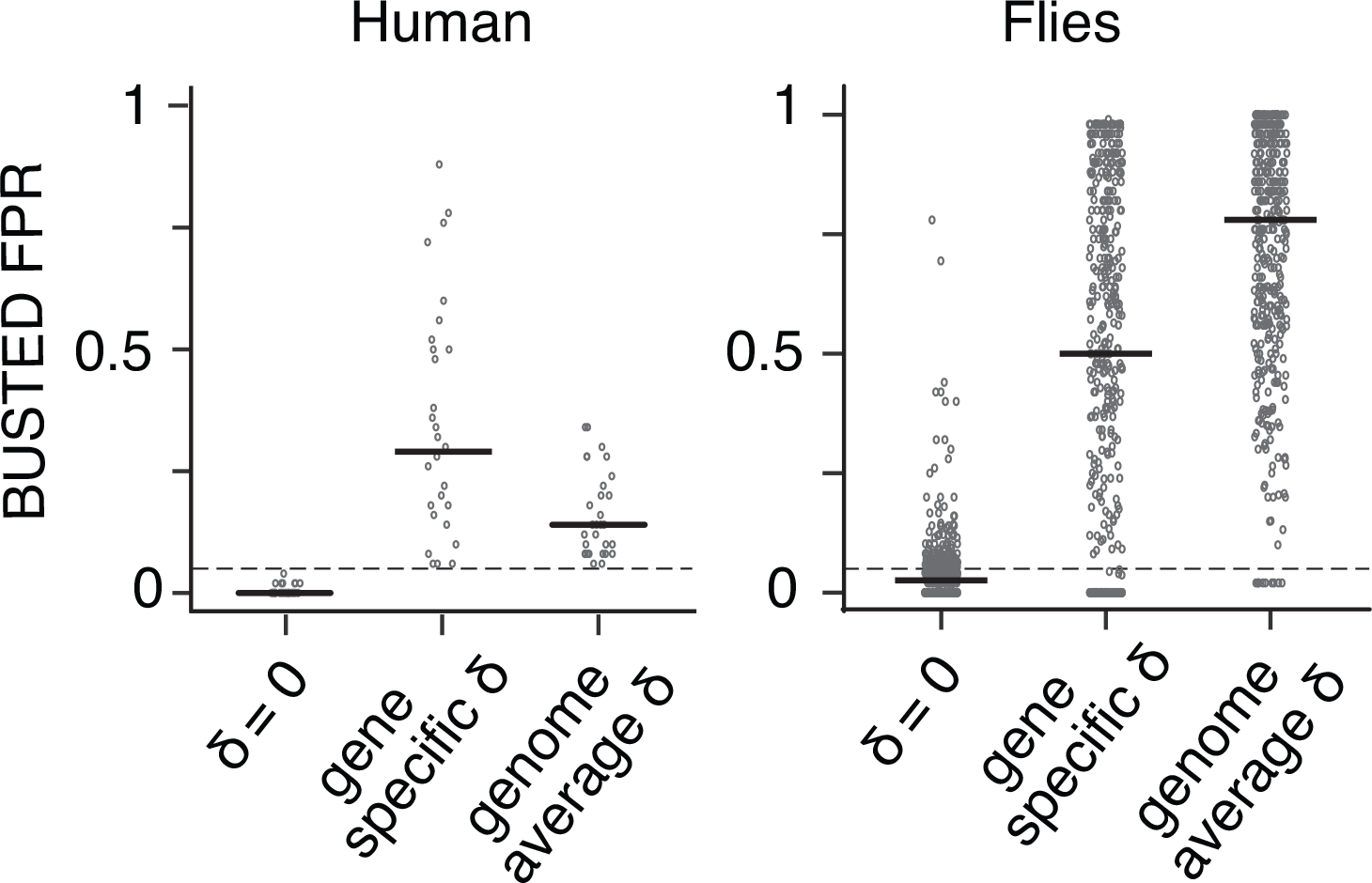
MNMs bias a newer test of positive selection. False positive inferences under realistic conditions using BUSTED. For every BST-significant gene in each dataset, 50 replicate alignments 5,000 codons long were simulated using the BS+MNM null model and parameter values estimated from the empirical sequences. These alignments were then analyzed for a signature of positive selection (P<0.05) using BUSTED. δ was assigned to its gene-specific estimate, to its average across all genes in each dataset, or to zero. FPR is the proportion of replicate alignments for each gene with P<0.05. Each dot represents the FPR for one gene; black bars are the median across genes.

### CMDs that invoke multiple nonsynonymous steps drive the signature of positive selection

Finally, we sought further insight into the reasons why CMDs yield a false signature of positive selection in the BST and related tests. In standard models of codon evolution, CMDs are interpreted as the result of two or more serial independent substitutions, even though they can be produced by MNMs in a single mutational event. We hypothesized that CMDs that imply multiple nonsynonymous nucleotide substitutions under these models would provide the strongest support for the positive selection model. We therefore classified CMDs in the empirical datasets by the minimum number of nonsynonymous single-nucleotide substitutions required from the ancestral to derived codon state under standard codon models. As predicted, we found that CMDs that imply more than one nonsynonymous step are dramatically enriched in BST-significant genes (**Fig 6a**).

**Figure 6.**
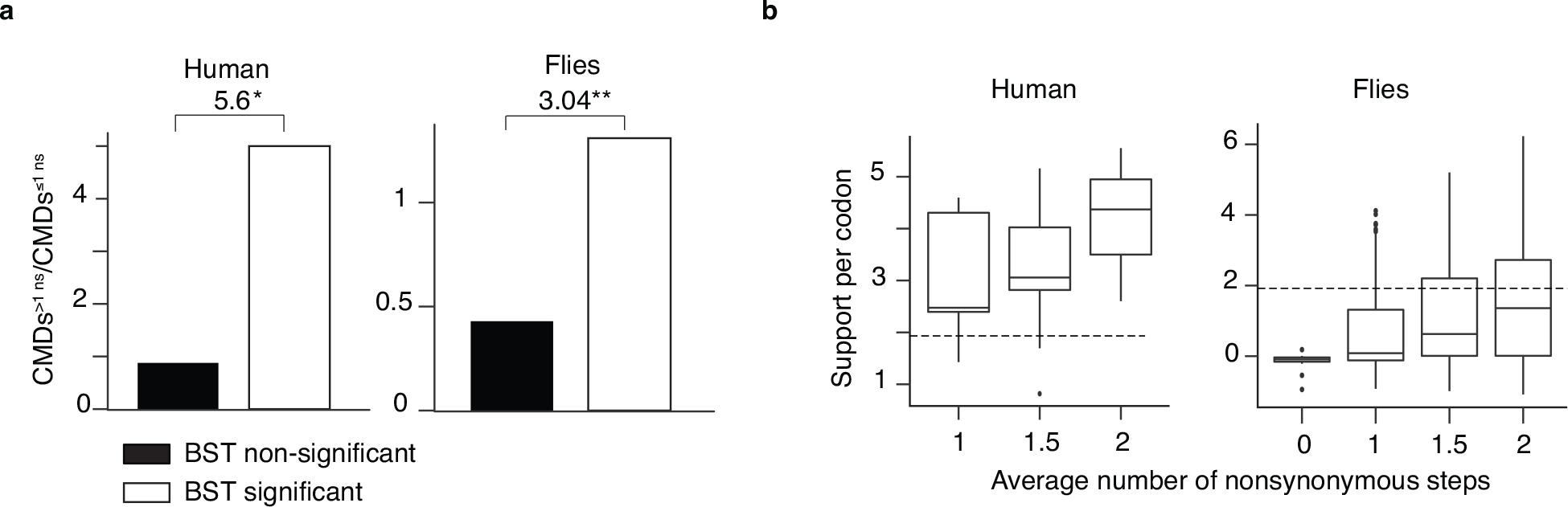
CMDs implying multiple nonsynonymous steps drive the BST. **(a)** For every CMD, the mean of the number of nonsynonymous single-nucleotide steps on the two direct paths between the ancestral and derived states was calculated. In BST-significant and BST-nonsignificant genes, the ratio of CMDs invoking more than one nonsynonymous step to those invoking one or fewer such steps is shown. Fold-enrichment is shown as the odds ratio. *, P=9e-04; **P= 1.6e-67 by Fisher’s exact test. **(b)** Support for the positive selection model provided by CMDs depends on the number of implied nonsynonymous single-nucleotide steps. Support is the log-likelihood difference between the positive selection and null models of the BST given the data at a single codon site. Box plots show the distribution of support by CMDs in BST significant genes categorized according to the mean number of implied nonsynonymous steps. Dotted line, support of 1.92, at which the BST yields a significant result for an entire gene (P<0.05). In human BST-significant genes, no CMDs imply zero non-synonymous changes.

We also examined the statistical support provided by different kinds of CMDs. As the number of nonsynonymous steps increased, the statistical support provided for the positive selection model also increased (**Fig. 6b**). CMDs that imply one nonsynonymous and one synonymous step typically provide weak to moderate support for the positive selection model, but CMDs that imply two nonsynonymous steps provide very strong support. In many cases, a single CMD in this latter category is sufficient to yield a statistically significant signature of positive selection.

## DISCUSSION

Our results demonstrate that the branch-site test suffers from a strong and systematic bias toward false positive inferences. This bias is caused by a mismatch between the method’s underlying codon model of evolution – which assumes that a codon with multiple differences can be produced only by two or more independent substitution events – and the recently discovered phenomenon of multinucleotide mutation, which produces such codons in a single event. Because of the structure of the genetic code and the high transversion rates that characterize MNMs, most codons produced by this mechanism cause more than one nonsynonymous single-nucleotide change. Confronted with this kind of codon data, the likelihood calculated by the BST is determined by the product of the probabilities of the individual mutations. Under the null model, the probability of such compound events is extremely small, but it can increase dramatically when *d*_N_/*d*_S_ exceeds one, as the positive selection model allows. This increase in likelihood afforded by the positive selection model is much greater than it would be if the substitution were interpreted as the result of a single multinucleotide event. Indeed, our results show that a single codon comprising two nonsynonymous substitutions is often sufficient to yield a statistically significant signature of positive selection in the BST for an entire gene.

As a result, CMDs are the primary drivers of positive results by the BST. Virtually all statistical support for positive selection in real alignments comes from CMD-containing sites; removing them from the alignment or incorporating MNMs into the BST’s model eliminates the signature of selection from the majority of genes. CMDs can be produced by either positive selection or by neutral evolution under multinucleotide mutation. In the former case, the BST will be correct; however, the test cannot reliably distinguish CMDs that represent authentic evidence of positive selection from those caused by MNM-induced bias.

The bias is strong and pervasive under realistic conditions. Indeed, when sequences were simulated under the null model using parameters estimated from the fly and mammalian datasets, the number of genes with false positive BSTs was approximately the same as the number of positive BST results when the empirical data were analyzed. There is therefore no excess of BST-positive results in these genomes beyond that potentially attributable to MNM-induced bias. Worse, these null simulations did not include the elevated transversion rate that characterizes MNMs, which exacerbates the test’s bias. Taken together, these results suggest the possibility that MNM-induced bias could explain many of the BST’s inferences of positive selection in these datasets.

Are our findings from these datasets generalizable? MNMs appear to be a property of all eukaryotic replication processes, and the MNM rates that we observed in mammals and flies are in the same range as those previously identified in genetic and molecular studies in a variety of eukaryotic species ^25,34,38^. Both datasets comprise a small number of taxa, but the BST seeks evidence of selection on individual branches, so it seems unlikely that larger trees will somehow inoculate the test against MNM-induced bias. We observed strong bias on lineages with divergence levels ranging from very low (on the human terminal branch) to moderate (the fly branches), so this problem does not appear to be unique to highly diverged sequences or phylogenies with long branches. We must therefore consider the possibility that many of the thousands of previously published reports of positive selection based on the BST could simply be the ones that happened by chance to neutrally fix one or more multinucleotide mutations.

We do not contend that the BST is always wrong or that molecular adaptive evolution does not occur. The existence of a bias, even a strong one, towards false positive inferences does not mean that all positive inferences are false: some of the CMDs in BST-significant genes may have evolved because of authentic positive selection, either by repeated substitution of single nucleotides in a codon or selection on MNMs. But because the BST test cannot distinguish the kinds of sequence data produced by positive selection from those produced by neutral evolution of MNMs, it does not provide reliable evidence that a gene evolved adaptively; nor does it provide a reliable estimate of the fraction of genes in a large set that evolved under positive selection. There are numerous cases of strongly supported adaptive evolution, such as those involving host-parasite and intracellular genetic conflicts, that have produced sequence signatures of positive selection in the BST and related tests that are likely to be authentic ^46^. The persuasive evidence in these cases, however, comes from sources other than the sequence signature.

If the BST and other lineage-specific tests based on the single-step codon model are unreliable in the face of multinucleotide mutation, what should researchers do? The BS+MNM test could be used to accommodate multinucleotide mutation; our results suggest this may be a promising approach. But there are many forms of evolutionary complexity that are not incorporated in this model, such as MNMs that affect three consecutive nucleotides in a codon, elevated transversion probability within MNMs, and many other kinds of heterogeneity that might bias the BS+MNM test ^47–49^. Other models are also available to incorporate MNMs ^9^, but their accuracy and robustness are not well characterized, either. More work is therefore required before the BS+MNM or similar models can be used with confidence in the branch-site or similar tests.

A complementary approach is to use functional experiments to explicitly test hypotheses that specific historical changes in molecular sequence caused changes in function or phenotype thought to have mediated adaptation ^50,51^. Indeed, the bias we observed may help to explain why some molecular experiments have shown that codons with a high posterior probability of positive selection in the BST do not contribute to putative adaptive functions, whereas the codon changes that do confer those functions have low or moderate PPs ^52^. Experimental tests provide the most convincing evidence of a gene’s putative adaptive history, but they require time-consuming laboratory and fieldwork ^53,54^, so it is not clear how to implement them on a genome-wide scale. Future research may develop and validate more robust models to detect positive selection, and these may help to identify candidate genes for which specific, testable hypotheses of past molecular adaptation on specific phylogenetic lineages can be formulated. The test primarily used for this purpose till now, however, is unreliable.

## ACKNOWLEDGEMENTS

We are grateful to the members of the Thornton lab for discussion and helpful comments. We thank the Beagle2, Midway2, and Tarbell supercomputing clusters at the University of Chicago. We also thank the developers of HyPhy for presenting an open source platform that allows limitless customization of standard analyses. Funding was provided by NIH R01GM104397 and R01GM121931 (JWT), NSF DEB-1601781 (JWT and AV), NSF DBI-1564611 (MWH), and the Precision Health Initiative of Indiana University (MWH).

## AUTHOR CONTRIBUTIONS

Analyses were designed by all authors, performed by AV, and interpreted by all authors. The manuscript was written by AV and JWT with contributions from MWH.

## METHODS

### Datasets, quality control, and inference of BST-significant genes

We analyzed two previously published comprehensive datasets of protein-coding alignments on a genomic scale, one in six mammals, the other in six *Drosophila* species **(Supplementary Table 2)** ^13,15,45^. We aimed to apply the branch-site test on every terminal lineage in the *Drosophila* dataset, and on the human lineage in the mammal dataset. We only retained gene alignments without gross misalignments, possessing complete coverage in all fly species, and minimally all primate species. We then applied the branch-site test as implemented in CODEML 4.7 to each alignment, assuming the phylogenetic relationships reported in the published studies (**Supplementary Fig. 2**) ^13,15^. Branch lengths and model parameters were estimated for each alignment by maximum likelihood (ML), and the F3×4 model was used for codon frequencies. We tested each gene in mammals for selection on the terminal branch leading to humans; in flies, each gene was tested separately for selection on each of the six terminal branches, and we express the fraction of positive inferences across genes as the proportion of all tests conducted ^6^. As is standard practice, we calculated P-values using a likelihood ratio test with 1 df (χ_1_^2^) which makes the test conservative under the null hypothesis ^6^. Genes were initially identified as having a putative BST signature of selection at P<0.05. We then applied a correction for multiple testing to a false discovery rate (FDR) <0.20 using the *q-value* package in R (available at http://github.com/jdstorey/qvalue).

To facilitate unambiguous analysis of CMDs, we removed genes containing CMDs falling in gaps. We also removed genes for which the ML ancestral reconstructions reported by CODEML at the base of the tested branch differed between the null and positive selection models, yielding a set of genes with CMDs that do not depend upon which model is chosen. In flies, 443 gene-tests (“genes”) were retained after these filters and constitute the BST-significant set of genes from this dataset. No genes on the human lineage were significant after FDR correction, so we retained as the BST-significant set from this dataset those genes that passed the ancestral reconstruction filter and had P<0.05 (**Supplementary Table 2**). The BST-nonsignificant set of genes comprises all genes that pass the alignment and ancestral reconstruction filter that are not in the BST-significant set (*n*=6757, humans; *n*=6883, flies). We also repeated our analysis of CMD enrichment (see below) using a gene set that had not been filtered for reconstruction consistency and found that our conclusions were unchanged (**Supplementary Table 7**)

We only considered genes where the ancestral codons (both CMD and non-CMD codons) have the same reconstruction under the BST null and BST alternate models. In doing so, we have also excluded CMDs in codons with gaps in the alignment. For example, in the human dataset, of the 82 genes that initially provided support for positive selection, 30 genes consist of unambiguously reconstructed codons under the null and alternate model (the BST-significant gene set). In 49 genes, CMDs fall in gaps. We did not consider the ancestral codon reconstructions at these sites, and excluded these from our analyses due to alignment ambiguities. The remaining 3 genes have CMDs that do not fall in gaps, for which the ancestral codons were reconstructed differently under the null and alternate models. If we re-consider these 3 ‘positively selected’ genes that were excluded, we find 3 additional CMDs, one in each of the genes. Including these genes made little to no difference to our CMD enrichment results.

### Support for positive selection

CMDs were identified in BST-significant and BST-nonsignificant genes as codons with 2 or 3 observed nucleotide differences between the ML states at the ancestral and extant nodes for the branch being tested; non-CMDs are codons with 0 or 1 differences on the branch tested. CMDs were not assessed on branches not tested.

To determine the role of CMDs in significant results from the BST, we excluded codon positions in BST-significant genes containing CMDs, reanalyzed the data using the BST, and calculated the fraction of tests that retained a significant result (P<0.05).

We quantified the proportion of statistical support for positive selection in BST-significant genes that comes from CMDs as follows. The site-specific support provided by one codon site in an alignment is the difference between the log-likelihoods of the positive selection model and the null model given the data at that site. Support for positive selection provided by all CMDs in a gene (*support_CMD_*) is the support summed over all CMD sites in the alignment. The proportion of support provided by CMDs is *support_CMD_ / (support_CMD_* + *support_nonCMD_)*. This proportion can be greater than 1 if support by non-CMDs is negative, as occurs if the likelihood of the null model at non-CMD sites is higher than that of the positive selection model, given the parameters of each model estimated by ML over all sites.

Sites were classified *a posteriori* as under positive selection if their Bayes Empirical Bayes posterior probability of being in class 2 (ω_2_>1) under the positive selection model in CODEML was >0.5 (moderate support) or >0.9 (strong support).

We categorized observed CMDs by the minimum number of nonsynonymous single-nucleotide steps implied under the Goldman-Yang model between the ancestral and derived states. For each CMD comprising two nucleotide differences, there are two paths by which they can be interconverted by two single nucleotide steps. We determined whether the steps on these paths would be nonsynonymous or synonymous using the standard genetic code and then calculated the mean number of nonsynonymous steps averaged over the two paths. Paths involving stop-codons were not included. We conducted a similar analysis for all possible CMDs in the universal genetic code table.

### BS+MNM codon substitution model and test

The codon substitution model of the classic BST is based on the Goldman-Yang (GY) model ^5^. Sequence evolution is modeled as a Markov process, where the matrix element *q_ij_*, the instantaneous rate of change from ancestral codon *i* to derived codon *j*, is defined for four types of changes: synonymous transitions and transversions, and nonsynonymous transitions and transversions (see *q_ij_* equation 1). Three parameters are estimated from the data by maximum-likelihood: ω, the ratio of nonsynonymous substitution rate to the synonymous substitution rate (*d*_N_/*d*_S_); π_j_, the equilibrium frequency of codon *j;* and κ the transversion:transition rate ratio.

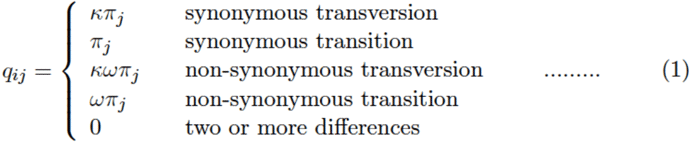

Element *q_ij_* is zero for substitutions involving more than one difference, so codons with multiple differences can only evolve through intermediate codons that are a single change away. A scaling factor applied to the matrix ensures that branch lengths are interpreted as the expected number of substitutions per codon.

We developed a modification of the GY model that incorporates MNMs using the parameter, δ, which represents the relative instantaneous rate of double substitutions to that of single substitutions (see *q_ij_* equation 2). When δ = 0, the BS+MNM model reduces to the classic BST model that does not incorporate MNMs (*q_ij_* equation 1). Triple substitutions have an instantaneous rate of zero.

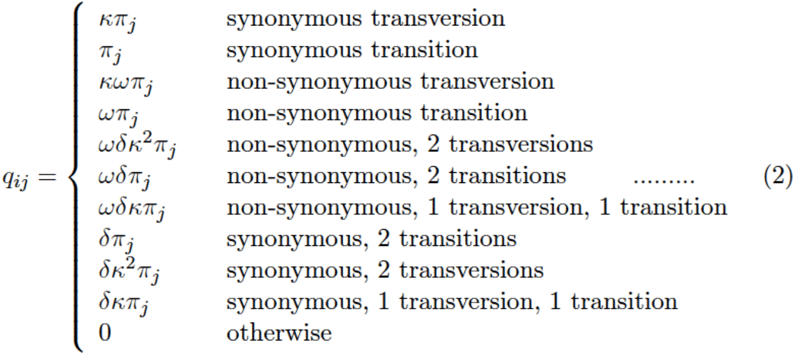

The BS+MNM test of positive selection is identical to the BST, except it utilizes this MNM codon model. We implemented this test by modifying the branch-site test batch file (YangNielsenBranchSite2005.bf) in Hyphy 2.2.6 software by declaring δ a global variable, incorporating it into the codon table, and allowing it to be optimized by ML as it other model parameters are.

We validated the BS+MNM implementation by simulating 50 replicate alignments using the BS+MNM null model in Hyphy under genome-median parameters (see below). We then used the BS+MNM procedure to find the ML estimate of each parameter, including branch lengths, given each alignment and the topology of the phylogeny used to generate the sequences. We compared the distribution of estimates over replicates to the “true” values used to generate the sequences (**Supplementary Fig. 3**).

To test if there is statistical support in the data for the BS+MNM null model relative to the standard BST null model, we performed an LRT with 1 df, comparing the fit of the BS+MNM null model and the BST null model on our empirical genes. Briefly, for each of the 6868 human genes, we tested if the BS+MNM null model fit the data better than the BST null model at P<0.05 and also applied and adjustment for multiple testing (FDR<0.2). We performed similar LRTs for each of the six terminal lineages in flies. To determine whether this test might be prone to falsely infer support for the BS+MNM model, we simulated control sequences under the null BST model with parameters derived from the empirical sequences and performed the LRT as described above. Only 2 percent of genes in humans and 2.6 percent in flies yielded significant support for BS+MNM at P < 0.05. Zero human genes and 0.006 percent of fly genes retained significance after multiple testing adjustment (FDR <0.2). (**Supplementary Table 4**).

### Simulations and analysis of false-positive bias

To characterize bias in the BST and other tests of selection, we conducted sequence simulations in the absence of positive selection under empirically derived conditions. We used the BS+MNM method we implemented in Hyphy to estimate by maximum likelihood (ML) the gene-specific branch lengths and parameters of the null BS+MNM model for every gene in the mammalian and fly datasets. We also calculated the genome-wide median of each parameter over all genes in each dataset (the “genome-average” parameter value). Probability density characterizations for parameters δ and gene length were performed using the *density* function in R.

We simulated sequence evolution under the BS+MNM null model using either gene-specific or genome-median parameters. First, we simulated a “pseudo-genome” without positive selection by simulating one replicate of each of the 6868 and 8564 mammalian and fly alignment, each at its empirical length, using the BS+MNM null model and the ML parameter estimates inferred for that gene from the empirical data. We then ran the BST on these sequences, testing for signatures of positive selection on the human lineage and each terminal fly lineage (**Supplementary Table 2**). Control simulations were conducted under identical conditions but with δ=0.

To test the effect of gene length on bias in the BST, we focused on genes in the BST-significant set. For each gene’s gene-specific parameters, we simulated 50 replicates alignments of length 5,000 or 10,000 codons. We analyzed these alignments using the BST, assigning the human branch as foreground for mammalian genes or, for flies, the same branch that produced a significant result when the empirical data were analyzed. The false positive rate (FPR) for any gene’s parameters is the fraction of replicates yielding a positive test (P<0.05). We also repeated these simulations and analyses using the genome-median value of δ. For control experiments without MNMs, we set δ =0 in the simulations.

To test the effect of the rate at which MNM substitutions are produced on false positive inference rates, we simulated evolution of alignments 5,000 codons long under the BS+MNM null model, using genome-median estimates for all parameters except δ, which we varied. At each value of δ, we simulated 50 replicates. We analyzed each replicate using the BST for selection on the human or *D. simulans* lineages and calculated the proportion of replicates for each value of δ that yielded a false positive inference (P<0.05).

We computed the observed proportion of tandem substitutions as a fraction of all substitutions on the human and *D. melanogaster* lineages in both empirical and simulated datasets. For each of the 6868 genes in the curated mammalian dataset, we aligned the human gene to the inferred sequence of the human-chimp ancestor, identified all substitutions as differences between these sequences, and calculated the proportion of tandem substitutions, T, as the number of substitutions at adjacent sites divided by the sum of substitutions at adjacent sites and those at non-adjacent sites across all sites in the dataset. Differences at adjacent sites were counted as a single tandem substitution. For each of the 8564 genes in the fly dataset, we aligned the *D. melanogaster* sequence to the *D. melanogaster/D. simulans* ancestor and followed the procedure described above. For simulated sequences, we repeated this procedure using the sequences simulated under the BS+MNM null model and parameters estimated from each gene in the empirical datasets.

### BUSTED

To examine the accuracy of BUSTED, we used Hyphy software 2.2.6 (batch files BUSTED.bf and QuickSelectionDetection.bf). We analyzed the 5,000 codon-long alignments simulated under the BS+MNM null model, using parameters estimated by ML for each BST-significant gene, with δ assigned either to its gene-specific estimate, its genome-average, or to zero. We applied BUSTED to the replicate alignments to test for selection (P<0.05) on the human lineage or the same fly lineage that was significant for that gene in the BST of the empirical data.

### Power analyses

To characterize the statistical power of the BST and BS+MNM tests, we simulated sequence evolution with positive selection of variable intensity and pervasiveness (**Supplementary Fig. 4**). Specifically, we used the BS positive model in Hyphy to simulate sequence evolution with the human and *D. simulans* terminal branches as the foreground branches. We used genome-average estimates of all parameters, including gene length (418 and 510 codons for mammals and flies, respectively), but we varied ω_2_ and p_2_. 20 replicate alignments were simulated under each set of conditions and then analyzed using the BST, the BS+MNM test, or BUSTED. For each set of conditions, the true positive rate was calculated as the fraction of replicates yielding a significant test of positive selection (P<0.05 for BST and BS+MNM, FDR<0.20 for at least one site in the alignment for BUSTED).

### BS+MNM+ κ_2_ model

We developed the BS+MNM+ κ_2_ model, which incorporates into the BS+MNM model (*q_ij_* equation 2) two different transversion:transition rate ratio parameters, κ_1_ for single-site substitutions and κ_2_ for MNMs (see *q_ij_* equation 3). All free parameters of the model are estimated by ML given a sequence alignment. This model was implemented by further modifying our BS+MNM batchfile in Hyphy 2.2.6 software by declaring κ_2_ a global variable, incorporating it into the codon table, and allowing it to be optimized by ML as other parameters are in the batch file.

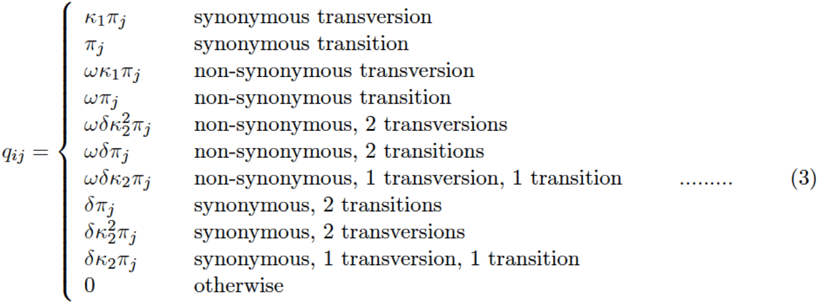

For validation, we estimated the parameters of the BS+MNM+ κ_2_ null model by ML for every alignment in each dataset and calculated the genome-average median estimate of each parameter (Fig. S7). We then simulated 50 replicate alignments of length 418 and 510 codons in the mammalian and fly datasets respectively, under the BS+MNM+ κ_2_ null model with all model parameters set to their genome-wide median. We then estimated each parameter by ML under the null model given each alignment and compared the distribution of estimates to the parameters used to generate the alignments. We found that most parameters were estimated accurately, but estimates of κ_2_ had high variance (**Supplementary Fig. S7**), presumably because the quantity of data in a single gene, in which CMDs are typically rare, is inadequate to support a robust estimate of this parameter. We therefore limited our use of this model to generating sequences by simulation rather than making inferences from sequence data.

To determine the effect of the MNM-specific transversion:transition rate on false-positive bias in the BST, we simulated sequences 10,000 codons long under the BS+MNM+κ_2_ null model, using genome-median parameters except κ_2_, which we varied. For each value of κ_2_, we simulated 50 replicates, applied the BST, and calculated the FPR as the fraction of replicates yielding a positive inference (P<0.05).

### Data availability

The empirical alignments reanalyzed in this study are available in the supplementary information of the original publications that generated these data ^12, 16, 45^.

### Code availability

The custom HYPHY batch codes for the BS+MNM and BS+MNM+κ_2_ tests are available as supplementary files and at https://github.com/JoeThorntonLab/MNM_SelectionTests.

